# Non-predictive online spatial coding in the posterior parietal cortex when aiming ahead for catching

**DOI:** 10.1101/197160

**Authors:** Sinéad A. Reid, Joost C. Dessing

**Affiliations:** School of Psychology, Queen’s University Belfast, David Keir Building, 18-30 Malone Road, BT9 5BN Belfast, Northern Ireland

## Abstract

Catching movements must be aimed ahead of the moving ball, which may require predictions of when and where to catch. Here, using Transcranial Magnetic Stimulation we show for the first time that, although interception movements were clearly aimed at the predicted final target position, the Superior Parietal Occipital Cortex (SPOC) displayed non-predictive online spatial coding. The ability to aim ahead for catching must thus arise downstream within the parietofrontal network for reaching.

---

The ability to predict future events in the environment is key to success of aspects of life. Some even argue that prediction is the key function of the brain^1^. Motor control is thought to rely heavily on prediction, because of sensorimotor delays and the general dynamic nature of the environment^2-4^. For instance, interceptive movements such as catching are typically aimed ahead of moving targets to account for any target displacement during movement execution^5^. This may involve predictions of where and when to intercept the object^6, 7^ - informing where to aim the movement and how fast to move, respectively. Counterintuitively, the resulting predictive behaviour – movements aimed towards a predicted future position of the moving target – may also emerge from control mechanisms that do not involve explicit predictions^8^. It is thus imperative to examine the neural basis of interception, which surprisingly has not been done extensively before in the neuroscience literature. The posterior parietal cortex (PPC) is likely involved, given its role in the online visual control of reaching^9-11^. Within PPC, the Superior Parietal Occipital Cortex (SPOC) and medial Intraparietal Sulcus (mIPS) purportedly code the retinotopic reach goal position^12, 13^ and the hand-target vector^12, 14-15^. For reaches to moving targets, the key question is whether this spatial coding reflects the continuously changing target position (non-predictive coding) or its future position (predictive coding). Here, we combined hemisphere-specific online repetitive Transcranial Magnetic Stimulation (rTMS) and interception of target trajectories crossing visual fields to test whether PPC uses predictive or non-predictive coding of moving targets during manual interception (**Fig. 1a**).

**Figure 1.**
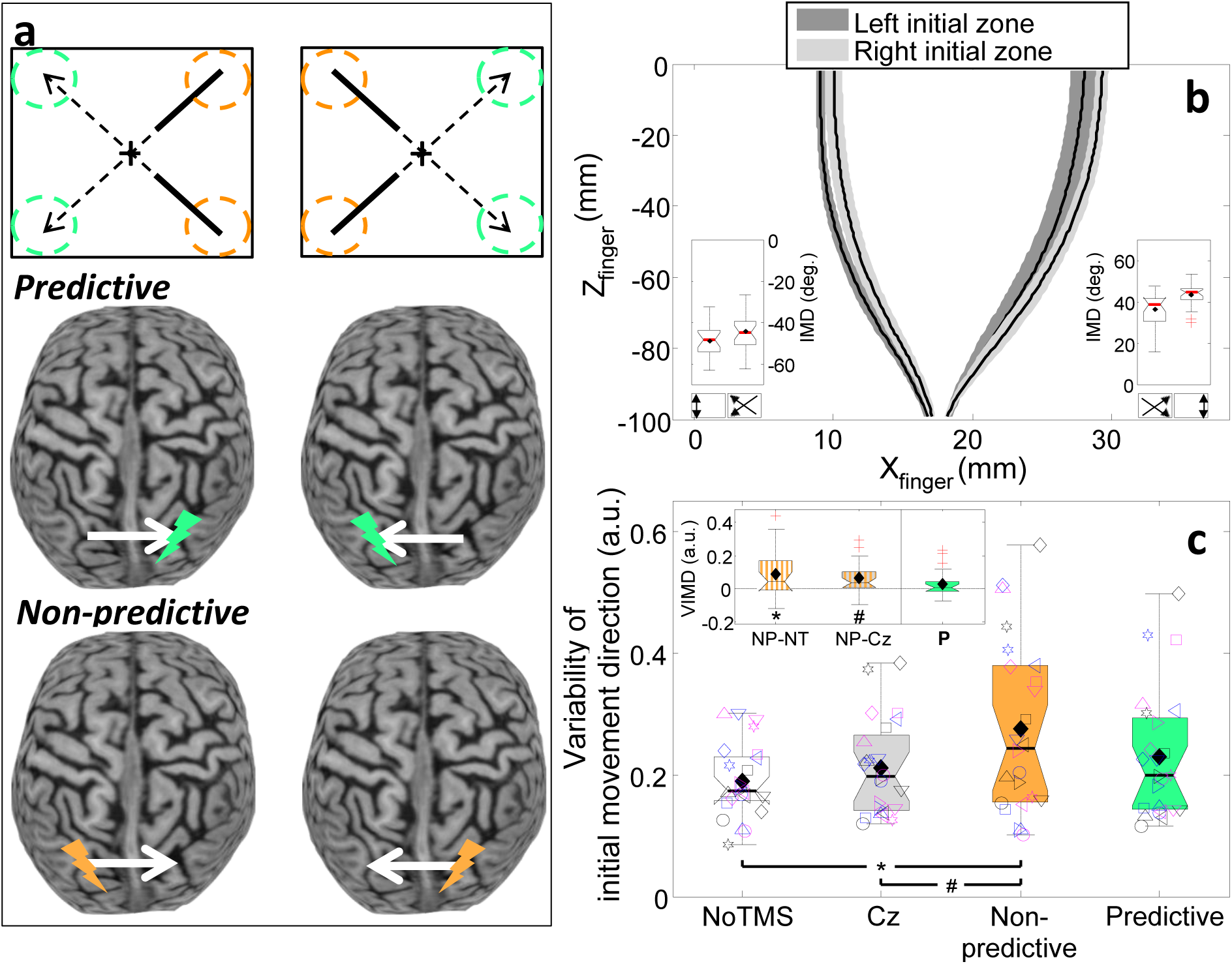
The rTMS paradigm and main findings for this study. **(a)** Predictive and Non-predictive retinotopic coding - concerning the final (green circles) and initial target position (orange circles), respectively - were tested using target trajectories crossing between visual hemifields (top; dashed for occluded part). Given central fixation (cross) and retinotopic coding in the PPC, effects are expected for rTMS to the hemisphere contralateral to the coded initial (orange lightning symbols) or final position (green lightning symbols). **(b)** Top view of a representative participant’s fingertip paths for conditions without TMS, averaged across vertical target motion directions (surface width represents the average s.d. across repetitions). The insets illustrate that across participants the initial movement direction (IMD) was in the direction of the final target zone; nevertheless, significant biases towards the initial target zone were also present (Left final zone: *χ*^2^(1) = 17.27, *p* = 3.2·10^−5^, Right final zone: *χ*^2^(1) = 23.68, *p* = 1.1·10^−6^; circular variant of *t*-test; α = 0.025). **(c)** Standard boxplot for the variability of the initial movement direction (VIMD) for rTMS to SPOC. The inset depicts the Non-predictive (NP) difference to the NoTMS (NT) and Cz conditions and the Predictive contrast (**P** = P-(NT+Cz)/2). Individual data is depicted using unique symbol/colour/jitter combinations. Means are indicated by black diamonds; outliers are red crosses. *: *t*(23) = 3.08, *p* = 0.0026, *d*’ = 0.63, α = 0.00625; #: *t*(23) = 3.26, *p* = 0.0017, *d*’ = 0.67, α = 0.00313; one-tailed paired-samples *t*-tests. Validity of results (*p*-values) was confirmed using 1,000,000 bootstraps.

In our experiment, participants reached to intercept a target moving down or up on a screen from either side to either side of the screen. Although the initial movement direction was significantly biased towards the initial target zone, movements were clearly planned ahead and aimed towards the final target zone (see **Fig. 1b**). We evaluated how these movements were affected by rTMS. We anticipated rTMS to yield an increase in movement variability, specifically of the initial movement direction for mIPS and horizontal interception error for SPOC. The rTMS effect was only significant for SPOC for the variability of initial movement direction and, most importantly, only when applied to the hemisphere that would code the retinotopic target position during stimulation (*t*(23) = 3.45; *p* = 0.0011; *d*’ = 0.70; see **Fig. 1c**). This demonstrates non-predictive spatial coding for manual interception in SPOC. The much smaller and non-significant effect for the spatial variability at interception (**Supplementary Fig. 1**) suggest that the rTMS effect was instantaneous and transient – disappearing after the rTMS train ended. In support, we found that the rTMS effect was present only for participants with an early initiation (**Fig. 2a**). With the exception of a single participant, the rTMS effect was only shown if initiation occurred >100ms before the last TMS pulse, that is, if the initial movement direction was determined within the rTMS window (**Fig. 2b**). Of the early initiators, only two clearly showed combined Non-predictive and Predictive effects (**Fig. 2c**), which may reflect effects of current visual target position in combination with cross-hemispheric predictive gain modulation^16^. None of test for the initial movement direction and horizontal interception error reached significance (**Supplementary Fig. 1**).

**Figure 2.**
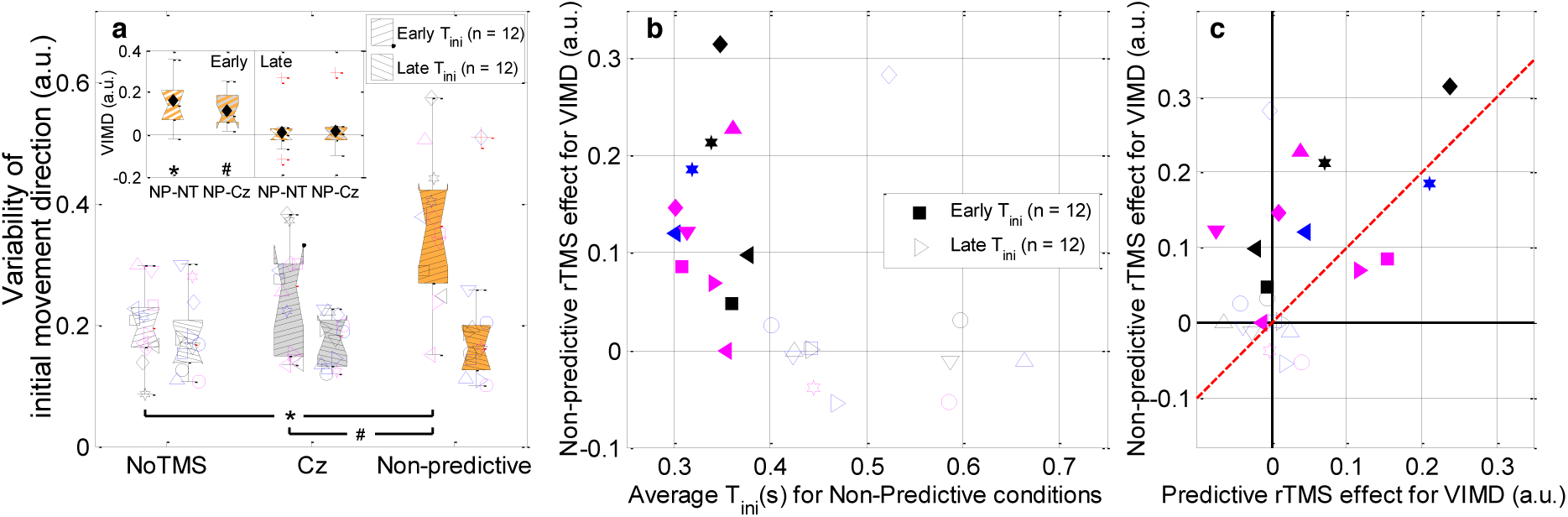
Group and individual differences in the non-predictive rTMS effects for SPOC on the variability of initial movement direction (VIMD). (**a**) Standard boxplot showing the effect broken down for two subsets of our sample, defined based on the moment of initiation (T_ini_). The inset depicts the differences between the Non-Predictive (NP) conditions and the NoTMS (NT) and Cz conditions for both groups. Individual data is depicted using unique symbol/colour/jitter combinations. Means are indicated by filled diamonds; outliers are red crosses. *: *t*(11) = 4.22; *p* = 0.00072; *d*’ = 0.86, α = 0.003125; #: *t*(11) = 5.04; *p* = 0.00019; *d*’ = 1.03, α = 0.00156. No significant differences were found for late initiators: NP-NT: *t*(11) = 0.37; *p* = 0.36; *d*’ = 0.076; NP-Cz: *t*(11) = 0.65; *p* = 0.26; *d*’ = 0.13. Validity of these one-tailed paired-samples *t*-tests (*p*-values) was confirmed using 1,000,000 bootstraps. (**b**) Individual Non-predictive rTMS effects (i.e., NP-(NT-Cz)/2) for SPOC on VIMD, as a function of the average T_ini_ in the Non-predictive conditions. With the exception of a single participant (open diamond), the rTMS effect was shown by participants initiating within 0.4s of target appearance. (**c**) Individual Non-predictive rTMS effects for SPOC on VIMD as a function of individual Predictive rTMS effects (i.e., P-(NT-Cz)/2) for SPOC on VIMD. The red dashed unity line indicates equal Predictive and Non-predictive effects. For most early initiators the rTMS effect predominantly occurred in the Non-predictive conditions; only two participants also showed considerable Predictive effects.

None of the rTMS effects on spatial movement features differed significantly between vertical target motion directions (all *p* > 0.10), nor between hemispheres receiving rTMS (all *p* > 0.20; see **Supplementary Tables 1 and 2**). Interestingly, the rTMS effects on spatial movement features were significant only for diagonal target trajectories that crossed visual hemifields; rTMS did not significantly affect movements towards targets that stayed within the same visual field (i.e., straight down or straight up) (all *p* > 0.022; see **Supplementary Table 3; Supplementary Fig. 2**). Indeed, for such same-field trajectories the contrast between SPOC and NoTMS/Cz was significantly smaller than the same contrast for diagonal trajectories (*t*(23) = 2.95; *p* = 0.0072; *d*’ = 0.60). Finally, no effects of rTMS to either SPOC or mIPS were observed on the temporal movement parameters (moment of initiation and movement time) (**Supplementary Figs. 1 and 2**).

The discussed effects of rTMS clearly demonstrate that SPOC uses non-predictive spatial coding for manual interception. The instantaneous and transient nature of these rTMS effects underscores PPC’s key role in online visual control of reaching^9-11^. Whereas studies involving stationary targets suggest SPOC codes the visual reach goal position^12, 13^ – the final target position in our task – this does not generalize to manual interception of moving targets. This demonstrates that, even if the brain forms an internal model of target motion^5^, SPOC does not simply read out a future target position from such a model to determine the reach goal when aiming ahead for manual interception.

Non-predictive online spatial coding within the PPC does not refute the use of predictive spatial control of manual interception. It does show that the ability to aim ahead of moving targets (**Fig. 1b**) must arise downstream from SPOC in the parietofrontal network. Although the rTMS effects at first sight might suggest SPOC codes current retinal target position, the significantly larger effect for targets crossing the vertical visual meridian (**Fig. 1c**, **Supplementary Fig. 2**) argues against this and may suggest a target motion dependence of the effect. Indeed, SPOC’s homologue in non-human primates (V6A) contains (non-predictive) spatially selective motion sensitive cells^17^. If SPOC’s activity effectively reflects target position and motion, future target positions may be decoded from its output. The predominant rTMS effect for diagonal target motion might suggest rTMS uniquely affected interhemispheric communication. Indirect support for this possibility comes from rTMS-induced disruptions of interhemispheric balance in PPC^18, 19^. A movement vector^14, 15^ based on the decoded predicted target position would be coded in the same hemisphere as the current target position (in SPOC) for non-diagonal target motion and in the opposite hemisphere for cross-meridian target motion (**Supplementary Fig. 3**). Whether the interception position is decoded, and thus explicitly represented, prior to or within the computation of the movement vector remains to be determined.

The aforementioned interpretation would be strengthened by the observation of Predictive rTMS effects for brain areas involved in the movement vector calculation. This calculation likely occurs within the PPC, anterior to SPOC^12, 14, 20^; we tested mIPS, but the absence of rTMS effects precludes any conclusions in this regard. Applying our paradigm (**Fig. 1a**) to other parts of the PPC likely involved in movement vector coding thus remains a key aim for future studies. More generally, understanding PPC’s movement-related activity during time-constrained reaching movements may prove seminal to creating effective neural prostheses that use PPC signals as input.

## Acknowledgements

We thank Fred Madelena and Kamil Kanas for their contributions to the experimental set-up and Rachel Giles for her help in running the experiments. The research leading to these results has received funding from the European Union Seventh Framework Programme FP7-CIG under grant agreement n° [334202], awarded to Joost C. Dessing, and from the Department of Education and Learning, Northern Ireland.

## Author Contributions

S.A.R. carried out all experiments, being accompanied by J.C.D during the first TMS sessions.S.A.R. and J.C.D designed the study and J.C.D implemented the experimental paradigm. J.C.D and S.A.R. conducted the data analyses and wrote the manuscript.

## Online Methods

### Subjects

Twenty-five participants took part in this experiment (mean age = 29, age range = 19-54, 13 females, 12 males). All had normal or corrected-to-normal vision and were right-handed (average laterality quotients: 92, range: 75-100^21^). Participants were recruited through social media advertisements and word of mouth. Participants completed two screening forms for contra-indications to MRI and TMS before participation was deemed safe. Participants also completed a fixation test to ensure they could comply with the fixation requirements of our task. In total, we recruited 44 participants; 19 participants were excluded because they failed the fixation pre-test. One participant only completed the SPOC session, while another participant’s data was excluded from the SPOC analyses due to an error in the TMS stimulator settings (5 instead of 6 pulses provided). As a result, 24 participants were included in both the SPOC and mIPS analyses, with 23 completing both sessions. Note that the exclusion of the participant from the SPOC analyses broke the perfect counterbalancing of rTMS conditions. Participants provided written informed consent before each session and completed an acute screening form before rTMS each session^22^.

### Experimental Set-up

Participants were seated in a height-adjustable chair behind a table on which a chinrest (with forehead support) and computer screen were mounted ∼35cm in front of the eyes (see **Fig. O1a**). Their head was fixed comfortably in the padded chinrest with a thick Velcro strap stretching across the back of the head to restrict excessive head movement. The experiment took place in a dark room; the only light sources were the stimuli presented on a 19inch Dell CRT screen (1280 x 1024 pixels, 75Hz refresh rate), a dim glow from the neuronavigation feedback screen and a small lamp that was switched on between blocks of trials. The light output of the CRT screen was reduced by two layers of darkening film (Defender Auto Window Film, Car Accessories Ltd., Buckingham, UK). Stimulus presentation was controlled through Matlab (The Mathworks, Nattick, MA, USA) by Version 3 of the Psychophysics Toolbox^23^. Movement of the right index finger was recorded with an NDI 3D Investigator system (Northern Digital Inc., Waterloo, ON, Canada) at 250Hz using six markers placed on an aluminium base taped to the distal part of the right index finger (partly covering the nail). This system was calibrated before each session such that the positive x-axis was rightward, the positive y-axis downward and the positive z-axis was into the screen (all from the participant’s perspective), with the origin in the left top corner of the screen. Binocular eye movements were recorded at 60Hz by SMI Eyetracking Glasses 2w Analysis Pro (SensoMotoric Instruments GmbH, Germany.

**Figure O1.**
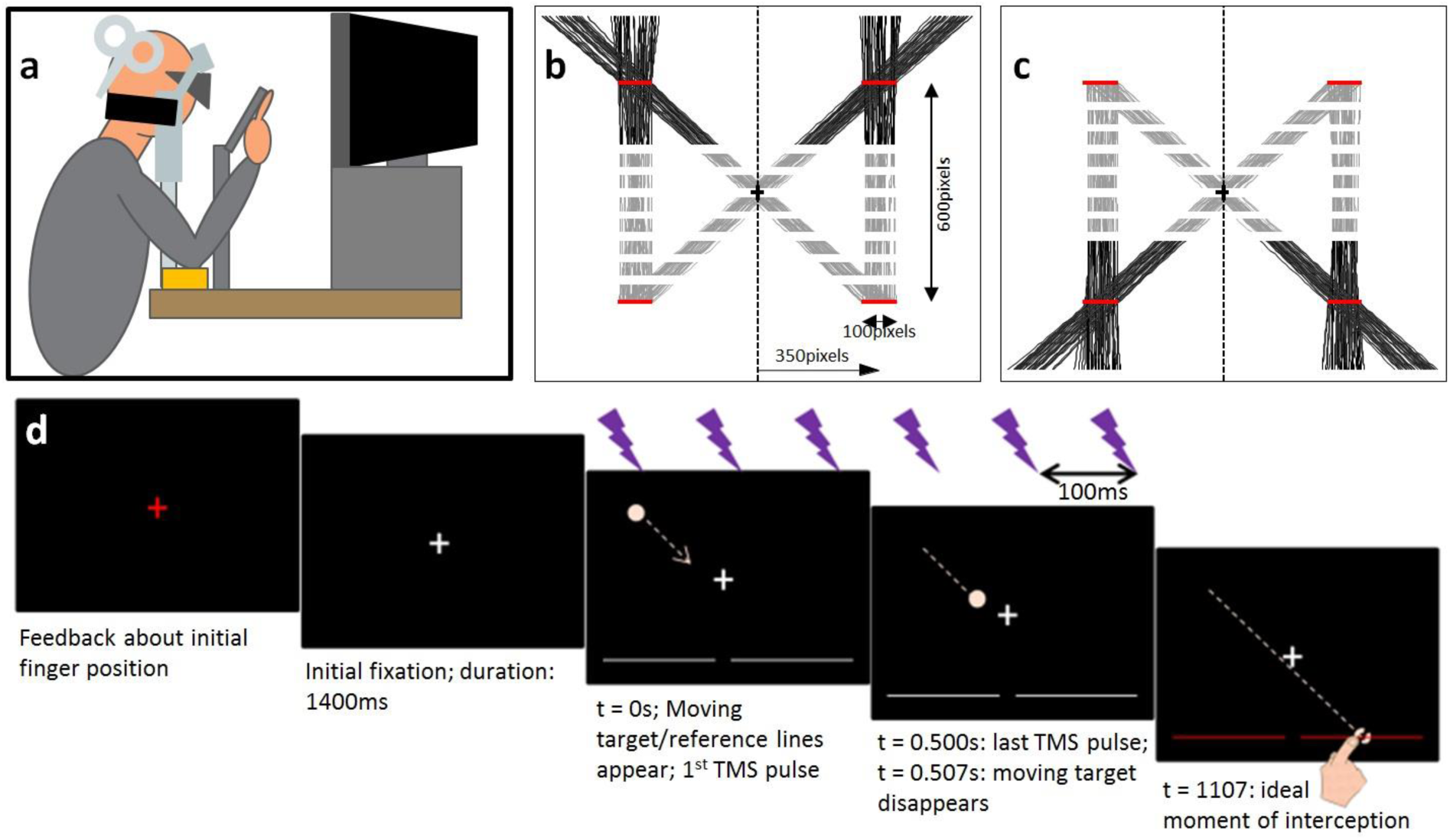
Experimental set-up and paradigm. **a)** Schematic side-view of the physical set-up. **b)** and **c)** Exemplary target paths used during the experiment for downward and upward target motion, respectively. Panels **b)** and **c)** represent the full screen at 1280 x 1024 pixels. The visible part of each path is shown as a solid red line; the occluded part is a grey dashed line. The initial and final target zones are displayed as thick horizontal red lines (not shown to participants). Note that the calibration involved pointing movements towards targets presented at the screen centre (black cross-sign) and the 8 outer positions of the initial and final target zones. Note that the initial zone constrained target positions midway through its visible window. **d)** Trial sequence. First, colour of the central fixation cross represents whether the fingertip is within (green) or outside (red) the initial finger zone. After the finger remained within the zone for 250ms, the white fixation cross is shown for 1400ms. Subsequently, the target appears (on TMS trials) in sync with the onset of the rTMS train (0-500ms, 6 pulses at 10Hz); two horizontal reference lines appear. After 38 frames (507ms) the target disappears, but it supposed to continue to keep moving until 600ms later (dashed circle), when the target reaches one of the reference lines, now turned red to signal the ideal moment of interception.

### Neuronavigation

To adequately position the TMS coil for each participant, we used a frameless stereotaxic neuronavigation system (Brainsight 2; Rogue Research, Montreal, Quebec, Canada). To this end, before the TMS sessions an MRI scan was obtained for each participant using a Siemens 1.5T scanner at NorthernMRI, Belfast (voxel size 1.0mm x 1.0156mm x 1.0156mm). We used six fiducial points (nasion, inion, right and left pre-auricular notch, right and left deepest point on the skull on the outside of the eye sockets) to align the MRIs with the recorded head position based on standard BrainSight methods. We deemed the alignment to be acceptable if the relative position of the fiducial points matches the positions obtained from the MRI with an error <2mm (typically ∼0.1-1mm).

We applied rTMS to the medial Intraparietal Sulcus (mIPS) and Superior Parietal Occipital Cortex (SPOC). mIPS was defined as a region located over the medial portion of the IPS, near the caudal part of the angular gyrus, in an attempt to match the sites used previously^14, 15^ (see **Supplementary Table 6**). SPOC was located on the superior part of the anterior bank of the parieto-occipital sulcus^12, 24^, (**see Supplementary Table 6**). A control site, Cz, was included to account for non-specific effects of rTMS and was defined as a point on the skull midway between the inion and nasion and equidistant from the left and right pre-auricular notches. The corresponding coil position was determined during practice stimulation at the start of the first session and used for neuronavigation during the session. During the experiment, feedback about the quality of neuronavigation was provided on a 15inch Dell LCD screen coated with several layers of the darkened film and with a carton cover on non-essential parts of the screen (to minimize the amount of light coming from this screen). Coil position was deemed acceptable if the distance from the optimal position on the skull (between the identified target spot and hotspot vector) was <1mm (typically 0.0-0.6mm).

### Target Trajectories

Target trajectories were varied in terms of vertical direction (upward or downward) and horizontal initial and final zone (see **Fig. O1b, c**). We thus presented four trajectory types: two approximately vertical trajectories that stayed within the same visual field, and two diagonal trajectories that crossed over from the left to the right visual field (or vice versa). The exact horizontal initial and final positions on a given trial were randomized within 100pixel wide zones centred on a horizontal eccentricity of ±350pixels relative to screen centre horizontal eccentricity (∼±16.2deg.; range 14.0-18.3deg.; see **Fig. O1b, c**). The ‘initial’ position was specified as the horizontal target position after 253ms (i.e., ∼midway through the TMS train). In combination with the occlusion time, this method ensured that the target disappeared well before crossing the midline. The vertical position of the zones was ±300pixels relative to the middle of the screen (depending on the direction of target motion) (∼±13.4deg. vertical eccentricity). Target trajectories were straight and of constant velocity.

### Procedures

Each session involved a calibration of the eye tracker (using a 3x3 grid spanning ±2.5 degrees in horizontal and vertical direction). The recorded raw gaze data for the outer eight points specified a zone (polygon) within which the gaze coordinates had to remain from target appearance until finger-screen contact (estimated online as the first time the fingertip came within 10mm of the screen surface) for the trial to be valid. The gaze data corresponded to the point of regard on the screen, averaged across the eyes; the experimenter could decide to ignore data from one eye if its calibration was deemed inadequate. Next, a calibration of the fingertip locations was performed. Participants reached to touch 9 points (i.e., screen centre and the edges of the horizontal range within which the final target positions could have been selected; see below) three times. The fingertip position relative to the six markers was determined based on the known physical location of the central point relative to these markers, averaged across the three touches. The other eight calibration points were used to reconstruct the ‘perfect’ final finger position (see Data Analyses).

The trial sequence is shown in **Fig. O1d**. With their heads fixed in the chin rest, participants positioned their fingertip within a pre-defined starting zone (determined using real-time streamed marker position data) between the eyes and the screen (100mm in front of and 10mm below the screen centre; spherical zone with radius of 15mm). During this phase, the fixation cross, presented at screen centre, was red while the fingertip was outside of the zone and green when it is inside. After the fingertip remained inside the starting zone for 250ms the fixation cross became white and was shown at the screen centre for 1400ms before a light pink target (∼6mm diameter) appeared either the top or bottom of the screen, moving at a constant speed to one of two unseen final target zones at the bottom/top of the screen. The full target trajectory took 1107ms (83 frames), the targets were occluded after 507ms visibility (38 frames). During the trial, participants moved their index finger from the initial position to reach and touch the invisible target as it reached the white line; to promote correct timing, the white line turned red when the invisible target reached it.

After each trial, gaze data was drift-corrected on each trial based on the median coordinates recorded in the last 300ms prior to target appearance. Eye tracking artefacts - infrequent sudden jumps (defined as gaze displacement during consecutive samples in opposite direction of at least twice the average amplitude of the saccade from screen centre to the four calibration points along the horizontal or vertical axes) - were removed. Any resulting gaps in the data were filled using linear interpolation. We recycled trials (at a random position in the remainder of the block) if fixation was inadequate (1256 trials, 8.0% of the original total number of trials), if reach initiation within 200ms of target appearance (19 trials, 8.0%), if initiation was not detected (defined at this stage as a >15mm displacement of the fingertip, relative to the position during the frame prior to the fixation point turning white) (in total 81 trials 0.5%), if fingertip-screen contact was not detected (120 trials, 0.8%), and if any interframe interval of our CRT screen deviated more than 20% from the optimal value (1/75) while the target was visible (185 trials, 1.2%). Since in many trials more than one criteria were met, a total of 1556 trials were rerun (10.0%). After trial parameters were saved, the next trial started.

Each participant completed a fixation test (10 practice trials followed by two blocks of 20 trials [with randomly selected target trajectories, see above]). At the end of this session, which took around 20 minutes to complete, the experimenter checked whether fixation was adequate in at least 30 of the 40 valid trials. This participant inclusion test ensured we only obtained an MRI scan for and applied rTMS to participants who could adequately perform our task. Included participants were taken to NorthernMRI to obtain their MRI scan and they completed two rTMS sessions, both of which contained blocks with TMS provided to Cz (a control site, included to control for non-specific aspects of the rTMS) and blocks without rTMS (to assess baseline behavioural performance). In one session additional blocks involved rTMS to left and right SPOC, while the other also involved TMS to the left and right mIPS. At the start of each rTMS session we determined the resting motor threshold (rMT) for both hemispheres prior to the eye tracker and fingertip calibrations and experimental blocks. This involved standard electromyography-based procedures (intensity/location for which 3 out of 6 single-pulse trials lead to >50μV peak-to-peak motor evoked potential of the first dorsal interrosus muscle). The rTMS sessions were at least 3 days apart.

Each experimental session started with a block of 10 practice trials without TMS. Per TMS condition (No TMS, Cz, left SPOC/mIPS, right SPOC/mIPS), participants completed two blocks of 42 trials (2 practice trials plus 5 repetitions per trajectory [2 vertical directions x 2 horizontal initial zones x 2 horizontal final zones]), plus any recycled trials (see above). All 4 conditions in each session were presented in blocks, the order of which was fully counterbalanced across participants (except for the additional participant in for SPOC); the two blocks per TMS condition were distributed within each session using an A-B-C-D-D-C-B-A design to control for the effects of fatigue. If the left and right rMT could differed, the stimulation intensity for one Cz block matched the left SPOC/mIPS value and for the other it matched the right SPOC/mIPS value. In each TMS trial six TMS pulses (10Hz, 120% rMT) were applied from target appearance, which implied the last pulse occurred just before target disappearance. The inter-trial interval was adjusted for safety reasons such that the time between TMS trains on subsequent trials was at least 6 seconds^22^. TMS was not provided if movements were initiated prior to the first TMS pulse (i.e., these trials were recycled). Participants were given a 5-10 minute break between TMS blocks; this allowed for cooling of the coil using ice packs and avoided any residual effects of stimulation. Note that during the early sessions some breaks occurred in the midst of blocks when the coil overheated, when fans were used for cooling. In those cases, the remaining trials in the block were completed directly after the coil had cooled down. In addition, the gaze data streaming sometimes malfunctioned, resulting in a short hold up. In all, these mid-block hold ups occurred 76 times during 288 blocks. Whenever deemed necessary, the eye tracker and fingertip calibrations were rerun (e.g., if eye tracking glasses or finger markers were moved on the body). Participants and experimenters wore earplugs throughout blocks of trials to protect from the TMS noises. All procedures were approved by the Queen’s University Belfast, School of Psychology Research Ethics Committee (45-2013) before any data was collected.

Note that one participant had a rather high rTM for the right hemisphere, which resulted in the rTMS being rather uncomfortable when its value was used to set the Cz intensity (it sometimes resulted in twitches of the eye muscles). During both sessions we therefore set the Cz position 2cm posterior and we stimulated Cz at 120% of the rMT of the left hemisphere. Incidentally, a few other instances of discomfort were reported (discomfort under ear, in eye, teeth clattering, left hand twitches, see log on https://osf.io/n4yjr/).

### Data Analyses

Data analyses were conducted offline using Matlab. Movement initiation was defined as the sample (until the fingertip was displaced 5mm relative to the position at target appearance) at which the forward fingertip velocity first last passed through the threshold on 5% of the maximal forward velocity achieved from target appearance to trial offset. The interception position was defined as the fingertip position at finger-screen contact. Finger-screen contact was determined as the first sample after the fingertip arrived within 10mm of the screen at which the forward fingertip velocity dipped below the threshold of 10mm/s. Contact-dependent variables were not calculated if the fingertip position could not be reconstructed (i.e., missing markers) for the sample preceding contact as well as ≥9 of the last 50 samples before contact. If this velocity-based definition did not work, contact was defined as the last valid sample (after the fingertip arrived within 10mm of the screen, but before 250ms after reaching its maximal forward position) at which the forward fingertip position was less than 1mm into the screen (n.b., due to soft tissue and bend of the finger, fingertip positions into the screen were possible). Note that these definitions of initiation and contact deviated slightly from the pre-registered definitions (see https://osf.io/n4yjr); minor algorithm adjustments were needed – prior to calculating the dependent variables – due to suboptimal visibility of the finger markers for some participants.

We determined the constant interception error (CE) by subtracting the ‘perfect’ fingertip position from the interception position. This perfect fingertip position was determined from the fingertip calibration: for each zone we used linear interpolation (based on the pixel coordinates of the exact final target position) between the fingertip positions recorded for the outer edges of the zone (averaged across the three repetitions in the initial calibration; note that a calibration point was excluded if it was further than 20mm from both other two repetitions). A positive horizontal CE was defined in the direction away from screen centre. Variable interception error (VE_x_) was defined as the variance of the horizontal CE across all repetitions for each condition. We applied a 4^th^ root transformation to VE_x_ to afford parametric statistics^25^. The initial movement vector (IMD) was defined as the 2D vector (i.e., in a horizontal plane, based on the lateral and forward coordinates) between the fingertip positions at initiation and 100ms after initiation. The variability of the initial movement direction (VIMD) was defined using the length of the within-condition average of the normalized initial movement vectors (*R*_*av*_), which we transformed to a linear scale to afford parametric statistics^26^, using:

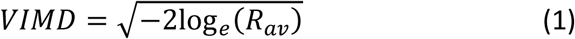

Due to marker occlusion and coil overheating, we excluded 71 trials from the CE_X_ and VE_X_ calculations (0.46%), 48 trials from the VIMD and IMD calculations (0.31%), and 43 trials from the calculation of the moment of initiation (0.28%), and 106 trials from the calculation of movement time (0.69%).

**Fig. 1a** illustrates our hypothesis tests, which follow from the retinotopic coding in mIPS and SPOC. Evidence for predictive and non-predictive coding was quantified using upward and downward diagonal target trajectories with stimulation to SPOC/mIPS in the hemisphere coding the visual hemifield of the final and initial target position, respectively. Dependent variables were averaged across these conditions; for Cz and NoTMS this did not require distinguishing between hemispheres. It should be noted that the hypothesis tests require that natural interception movements are planned ahead and not aimed at the current target position; this was visually confirmed (i.e., effects large enough not to require statistics); any additional (smaller) effects of the initial target zone on IMD was statistically tested separately for the two final target zones (circular test^26^, alpha = 0.025). Note that for this particular test a positive IMD denoted rightward (as opposed to the rTMS tests, where positive was in the direction of the final target zone).

For both PPC sites, we performed the comparisons based on a predicted increase in movement variability due to rTMS in comparison to both no TMS and Cz stimulation. Based on the literature, we anticipated and thus preregistered effects for SPOC on VEx^24^ and for mIPS on VIMD^14, 15^. In the preregistration, we treated the sessions (SPOC and mIPS) as separate experiments; starting at a Bonferroni-corrected alpha-level of 0.025 (i.e., 2 tests, referred to as ‘Predictive’ and ‘Non-predictive’ hereafter), we applied a further Holm-Bonferroni step-down correction for the comparisons with NoTMS and Cz.

We always anticipated further exploration of our data. Firstly, since we assumed rTMS would affect the movement from initiation to interception, the aforementioned tests should also show the effects on VIMD for SPOC stimulation and on VE_x_ for mIPS stimulation. A more conservative alpha-level was used for these tests; taking the exploration as part of testing rTMS effects for two hypotheses for two brain sites using two dependent variables, we started with an alpha-level of 0.05/8 = 0.00625. This alpha-level was applied to contrasts of predictive/non-predictive in this case compared to the average of NoTMS and Cz; only significant effects were followed up by separate comparisons with the NoTMS and Cz conditions with associated Holm-Sidak corrections as described above. Importantly, since we only expected rTMS-induced *increases* in movement variability all comparisons referred to above involved one-tailed paired-samples *t*-tests (with Cohen’s *d* reported for effect size).

The data was further explored using other dependent variables. We used the aforementioned statistical procedure to explore the effects for both PPC sites on IMD (in degrees) and the horizontal constant interception error (CE_x_ in mm); since we did not have a unique prediction about the direction of the effects these involved two-tailed paired-samples *t*-tests (n.b. a circular test was used for IMD^26^). Although the present investigation focused on spatial control, for completeness we also tested whether rTMS affected the moment of initiation (relative to target appearance) and movement time, both expressed in seconds.

For all tests, we explored whether there were any asymmetries in the rTMS effects between hemispheres and vertical target motion directions. We compared the size of the Predictive/Non-predictive contrasts (relative to the average of NoTMS and Cz) between vertical directions and between hemispheres (the same procedure for correcting alpha levels was used, except that no break-down for NoTMS and Cz was planned to avoid inflating the number of tests). We also evaluated the rTMS effects for non-diagonal trajectories for all dependent variables: we took the difference between the effects of target motion coded in the stimulated and non-stimulated hemisphere and contrasted this with the same differences for NoTMS and Cz (averaged across these two control conditions).

Based on one of the effects observed for rTMS to SPOC on VIMD, specific follow-up tests were designed. We compared the size of the Non-predictive contrast (relative to the average of NoTMS and Cz) to the same contrast for straight trajectories (coded within the stimulated hemisphere). We explored individual differences in the size of the Non-predictive and Predictive contrasts, to ascertain whether there was any indication of combined predictive and non-predictive effects^16^. Finally, we tested for Non-predictive coding separately for two groups, which differed in how soon they initiated their movements after target appearance. To this end, median-splitting based on the overall average moment of initiation for the Non-predictive conditions for SPOC was used to create groups). Validity of all statistical tests was confirmed using bootstrapped *p*-values (n = 1,000,000, see **Supplementary Tables 1-5**).

### Data Availability

All data and analysis code for this study are available for download through https://osf.io/n4yjr.

---

**Supplementary Figure 1.**
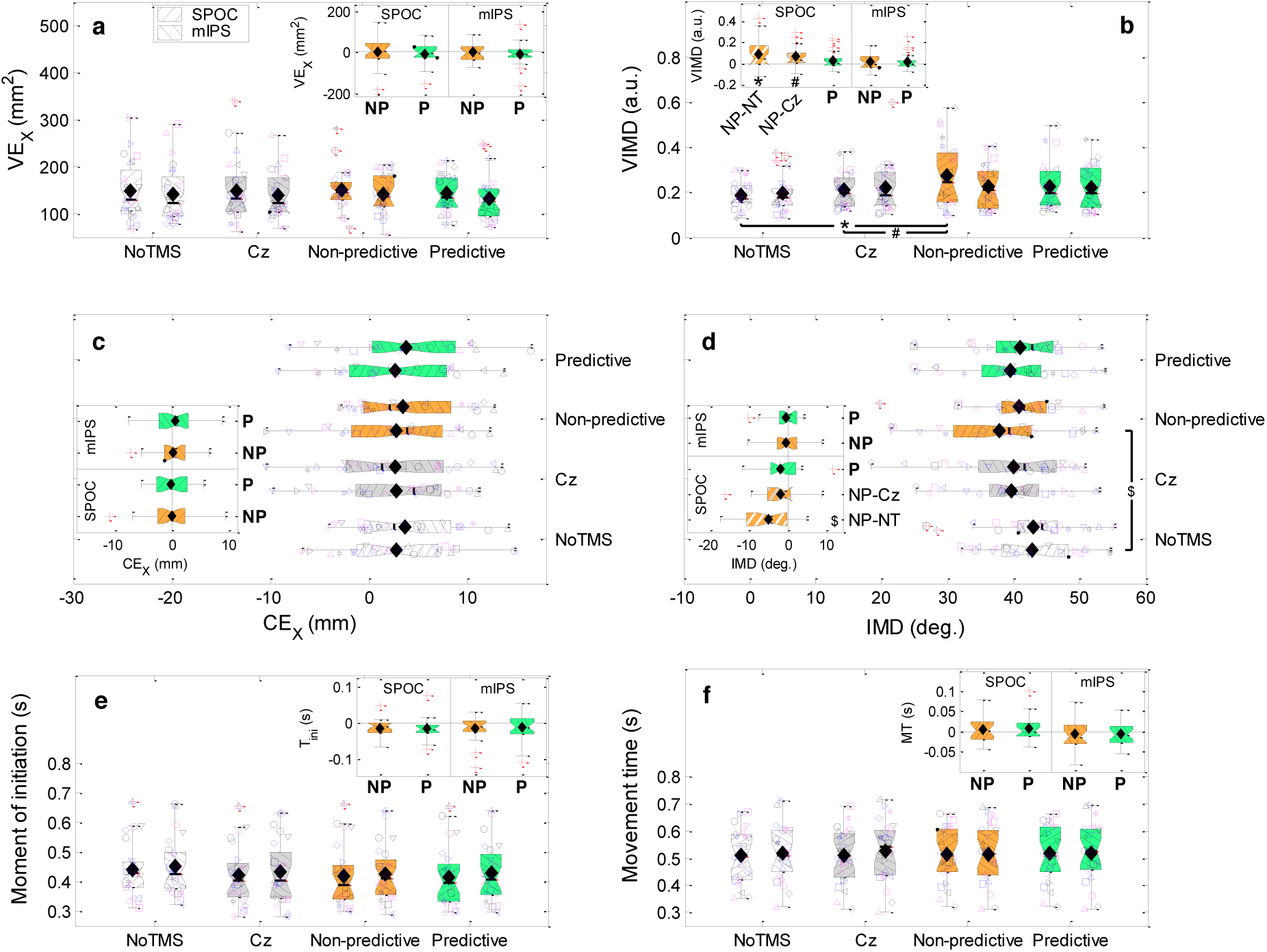
Standard boxplots for the Non-predictive and Predictive rTMS effects on all horizontal movement variability and biases and timing for SPOC and mIPS. (**a**) Shows the effects on the variability of the horizontal interception error (VE_X_). (**b**) Shows the effects on the variability of the initial movement direction (VIMD). (**c**) Shows the effects on the horizontal interception error (CE_X_). (**d**) Shows the effects on the initial movement direction (IMD). (**e**) Shows the effects on the moment of initiation (T_ini_). (**f**) Shows the effects on the movement time (MT). In all panels, the Non-predictive (NP) and Predictive (P) rTMS effects are shown relative to rTMS to the control site Cz and the condition without rTMS (NoTMS, NT). Insets show the Non-predictive (**NP** = NP-(NT+Cz)/2) and Predictive (**P** = P-(NT+Cz)/2) contrasts; for significant contrasts the individual comparisons with NoTMS and Cz are shown instead. In the main panels, individual data is depicted using unique symbol/colour/jitter combinations. Means are indicated by filled diamonds; outliers are red crosses. *: *t*(23) = 3.08, *p* = 0.0026, *d*’ = 0.63, α = 0.00625; #: *t*(23) = 3.26, *p* = 0.0017, *d*’ = 0.67, α = 0.00313 (one-tailed paired-samples *t*-tests). $: *χ*^2^(1) = 12.77; *p* = 0.00035, α = 0.00313 (main contrast: *χ*^2^(1) = 8.78; *p* = 0.0031) (circular one-sample tests [relative to 0]^26^). Validity of results (*p*-values) confirmed using 1,000,000 bootstraps.

**Supplementary Figure 2.**
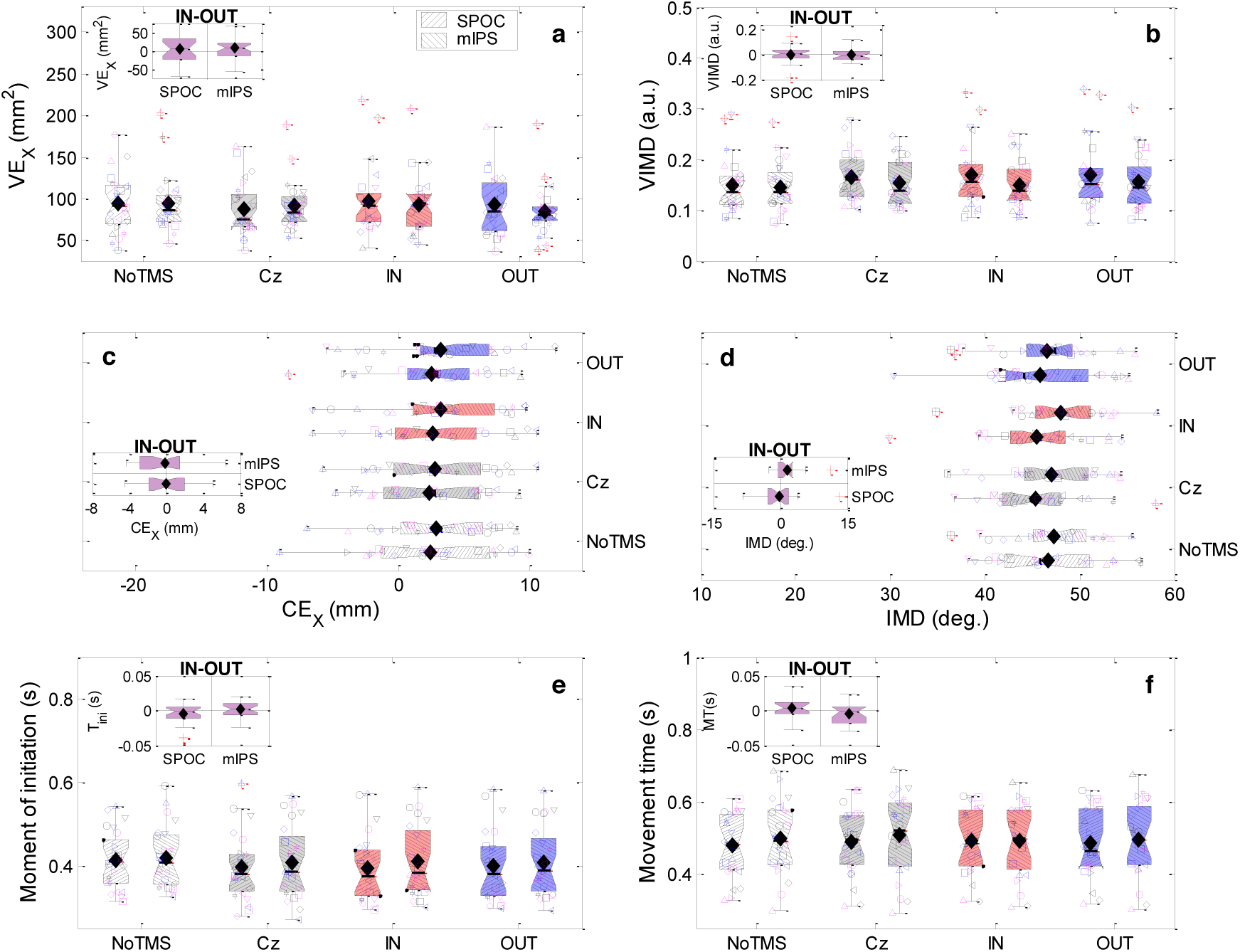
Standard boxplots for the rTMS effects for non-diagonal target trajectories for both SPOC and mIPS. (**a**) Effects for the variable error of the horizontal interception error (VE_X_). (**b**) Effects for the variability of initial movement direction (VIMD). (**c**) Effects for the horizontal interception error (CE_X_). (**d**) Effects for the initial movement direction (IMD). (**e**) Effects for the moment of initiation (T_ini_). (**f**) Effects for the movement time (MT). Insets show the contrasts between the rTMS effects for targets that, according to their retinotopic position, would be coded inside (IN) the stimulated hemisphere or in the opposite hemisphere (OUT). None of the differences were significant. Validity of results (*p*-values) confirmed using 1,000,000 bootstraps.

**Supplementary Figure 3.**
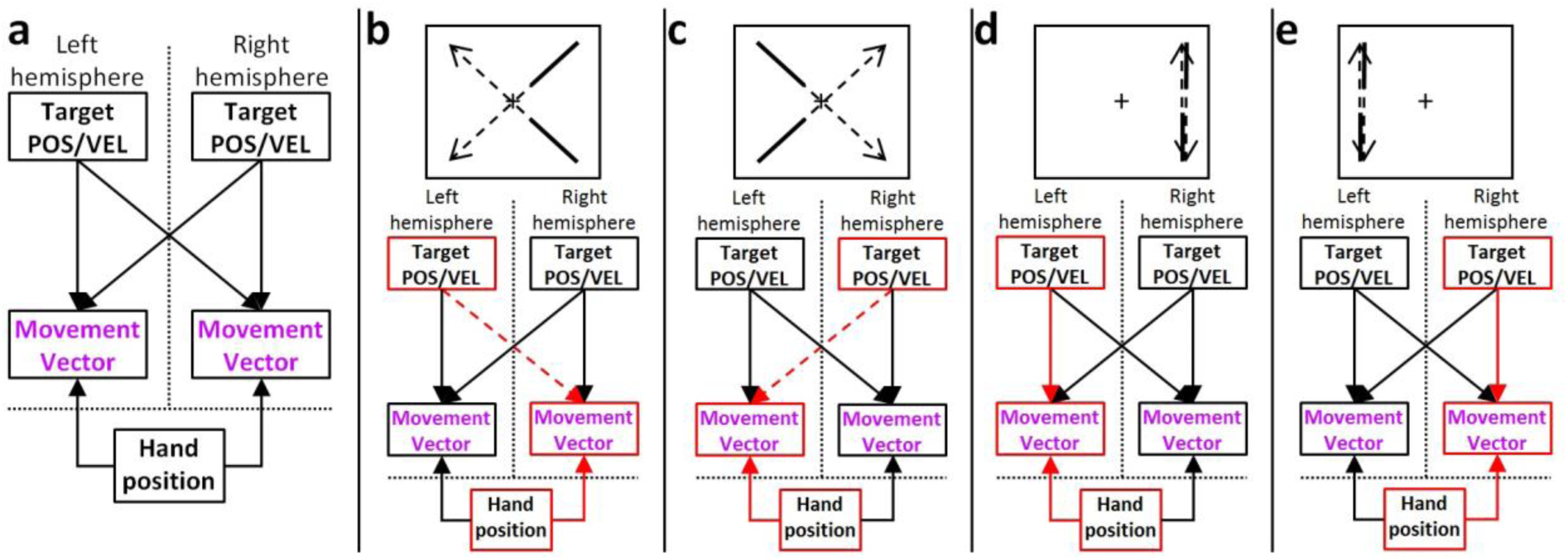
A schematic of hemisphere-specific movement planning. (**a**) Calculation of the movement vector in each hemisphere, based on combined target position and velocity signals (target POS/VEL) – originating in SPOC according to the interpretation of our findings – and hand position signals. Any hemisphere specificity of the latter is omitted because it is not pertinent to our interpretation. (**b**) The active nodes and pathways for diagonal target trajectories from the right to the left visual hemifield. (**c**) The active nodes and pathways for diagonal target trajectories from the left to the right visual hemifield. (**d**) The active nodes and pathways for non-diagonal target trajectories in the right visual hemifield. (**e**) The active nodes and pathways for non-diagonal target trajectories in the left visual hemifield. In all cases the target’s position and velocity are coded in the hemisphere opposite to the visual hemifield containing the current target position. For diagonal trajectories ((**b**,)&(**c**)), the movement vector is computed in the hemisphere opposite to the visual hemifield containing the *final* target position. For straight trajectories ((**d**)&(**e**)) the movement vector is coded in the hemisphere opposite to the visual hemifield containing the *current* target position. Our interpretation depends on rTMS predominantly affecting signals from SPOC that cross to the opposite hemisphere, which are displayed as dashed arrows in (**b**) and (**c**). For simplicity and lack of evidence, a stage explicitly coding the predicted future target position (using target position and velocity signals from SPOC) is omitted; our results do show that SPOC does *not* represent this stage.

**Supplementary Table 1:** Non-predictive and Predictive rTMS effects on spatial movement variability

| Site/DV | Test | Statistics | Bootstrapped $p$ |
| --- | --- | --- | --- |
| SPOC/$VE_x$ | Non-predictive | vs. NoTMS: $t(23) = 0.47$; $p = 0.32$; $d' = 0.096$ | $p = 0.32$ |
| | | vs. Cz: $t(23) = 0.48$; $p = 0.32$; $d' = 0.098$ | $p = 0.32$ |
| | Predictive | vs. NoTMS: $t(23) = -0.19$; $p = 0.57$; $d' = -0.039$ | $p = 0.58$ |
| | | vs. Cz: $t(23) = -0.14$; $p = 0.56$; $d' = -0.029$ | $p = 0.56$ |
| | (Non-predictive - (NoTMS+Cz)/2) UP vs. DWN | $t(23) = -1.28$; $p = 0.21$; $d' = -0.26$ | $p = 0.19$ |
| | (Predictive - (NoTMS+Cz)/2) UP vs. DWN | $t(23) = -0.30$; $p = 0.77$; $d' = -0.061$ | $p = 0.76$ |
| | (Non-predictive - (NoTMS+Cz)/2) L vs. R | $t(23) = -0.47$; $p = 0.65$; $d' = -0.095$ | $p = 0.64$ |
| | (Predictive - (NoTMS+Cz)/2) L vs. R | $t(23) = -0.0019$; $p > 0.99$; $d' = -0.00038$ | $p > 0.99$ |
| mIPS/$VE_x$ | Non-predictive | $t(23) = 0.32$; $p = 0.38$; $d' = 0.065$ | $p = 0.37$ |
| | Predictive | $t(23) = -0.71$; $p = 0.76$; $d' = -0.15$ | $p = 0.77$ |
| | (Non-predictive - (NoTMS+Cz)/2) UP vs. DWN | $t(23) = -0.26$; $p = 0.80$; $d' = -0.052$ | $p = 0.79$ |
| | (Predictive - (NoTMS+Cz)/2) UP vs. DWN | $t(23) = 0.49$; $p = 0.63$; $d' = 0.10$ | $p = 0.62$ |
| | (Non-predictive - (NoTMS+Cz)/2) L vs. R | $t(23) = 1.36$; $p = 0.19$; $d' = 0.28$ | $p = 0.16$ |
| | (Predictive - (NoTMS+Cz)/2) L vs. R | $t(23) = 0.75$; $p = 0.46$; $d' = 0.15$ | $p = 0.45$ |
| SPOC/VIMD | Non-predictive | **$t(23) = 3.45$; $p = 0.0011$; $d' = 0.70$**<br>**vs. NoTMS: $t(23) = 3.08$; $p = 0.0026$; $d' = 0.63$**<br>**vs. Cz: $t(23) = 3.26$; $p = 0.0017$; $d' = 0.65$** | **$p = 0.00045$**<br>**$p = 0.0018$**<br>**$p = 0.00092$** |
| | Predictive | $t(23) = 1.75$; $p = 0.046$; $d' = 0.36$ | $p = 0.044$ |
| | (Non-predictive - (NoTMS+Cz)/2) UP vs. DWN | $t(23) = 0.47$; $p = 0.64$; $d' = 0.096$ | $p = 0.63$ |
| | (Predictive - (NoTMS+Cz)/2) UP vs. DWN | $t(23) = -0.17$; $p = 0.87$; $d' = -0.035$ | $p = 0.86$ |
| | (Non-predictive - (NoTMS+Cz)/2) L vs. R | $t(23) = -0.20$; $p = 0.84$; $d' = -0.040$ | $p = 0.84$ |
| | (Predictive - (NoTMS+Cz)/2) L vs. R | $t(23) = 0.26$; $p = 0.80$; $d' = 0.052$ | $p = 0.80$ |
| mIPS/VIMD | Non-predictive | vs. NoTMS: $t(23) = 1.67$; $p = 0.055$; $d' = 0.34$<br>vs. Cz: $t(23) = 0.27$; $p = 0.39$; $d' = 0.056$ | $p = 0.047$<br>$p = 0.41$ |
| | Predictive | vs. NoTMS: $t(23) = -1.44$; $p = 0.082$; $d' = 0.29$<br>vs. Cz: $t(23) = 0.10$; $p = 0.46$; $d' = 0.021$ | $p = 0.076$<br>$p = 0.46$ |
| | (Non-predictive - (NoTMS+Cz)/2) UP vs. DWN | $t(23) = -1.12$; $p = 0.27$; $d' = -0.23$ | $p = 0.25$ |
| | (Predictive - (NoTMS+Cz)/2) UP vs. DWN | $t(23) = -0.0072$; $p = 0.99$; $d' = -0.0015$ | $p = 0.99$ |
| | (Non-predictive - (NoTMS+Cz)/2) L vs. R | $t(23) = 0.24$; $p = 0.82$; $d' = 0.048$ | $p = 0.81$ |
| | (Predictive - (NoTMS+Cz)/2) L vs. R | $t(23) = 0.43$; $p = 0.67$; $d' = 0.087$ | $p = 0.66$ |

Note: rTMS = repetitive Transcranial Magnetic Stimulation; DV = Dependent variable; SPOC = Superior Parietal Occipital Cortex; mIPS = Medial Intraparietal Sulcus; $VE_x$ = variance of horizontal interception error; VIMD: variability of initial movement direction; NoTMS = conditions without rTMS; Cz = conditions with rTMS applied to the control site Cz; UP = upward target motion; DWN = downward target motion; L = stimulation applied in the left hemisphere; R = stimulation applied in the right hemisphere.

**Supplementary Table 2:** Non-predictive and Predictive rTMS effects on spatial movement biases

| Site/DV | Test | Main contrast | Bootstrapped $p$ |
| --- | --- | --- | --- |
| *SPOC/CE<sub>x</sub>* | Non-predictive | $t(23) = -0.072; p = 0.94; d' = -0.015$ | $p = 0.94$ |
| | Predictive | $t(23) = -0.19; p = 0.85; d' = -0.04$ | $p = 0.84$ |
| | (Non-predictive - (NoTMS+Cz)/2) UP vs. DWN | $t(23) = 0.71; p = 0.49; d' = 0.14$ | $p = 0.47$ |
| | (Predictive - (NoTMS+Cz)/2) UP vs. DWN | $t(23) = -1.65; p = 0.11; d' = -0.34$ | $p = 0.091$ |
| | (Non-predictive - (NoTMS+Cz)/2) L vs. R | $t(23) = 0.80; p = 0.43; d' = 0.16$ | $p = 0.42$ |
| | (Predictive - (NoTMS+Cz)/2) L vs. R | $t(23) = -1.03; p = 0.32; d' = -0.21$ | $p = 0.29$ |
| *mIPS/CE<sub>x</sub>* | Non-predictive | $t(23) = 0.36; p = 0.72; d' = 0.074$ | $p = 0.71$ |
| | Predictive | $t(23) = 0.80; p = 0.43; d' = 0.16$ | $p = 0.41$ |
| | (Non-predictive - (NoTMS+Cz)/2) UP vs. DWN | $t(23) = 0.11; p = 0.92; d' = 0.022$ | $p = 0.91$ |
| | (Predictive - (NoTMS+Cz)/2) UP vs. DWN | $t(23) = -1.61; p = 0.12; d' = -0.33$ | $p = 0.099$ |
| | (Non-predictive - (NoTMS+Cz)/2) L vs. R | $t(23) = 0.14; p = 0.89; d' = 0.029$ | $p = 0.88$ |
| | (Predictive - (NoTMS+Cz)/2) L vs. R | $t(23) = 0.54; p = 0.60; d' = 0.11$ | $p = 0.58$ |
| *SPOC/IMD* | Non-predictive | $\chi^2(1) = 8.78; p = 0.0031$<br>vs. NoTMS: $\chi^2(1) = 12.77; p = 0.00035$<br>vs. Cz: $\chi^2(1) = 2.61; p = 0.11$ | $p = 0.00067$<br>$p = 0.000017$<br>$p = 0.086$ |
| | Predictive | $\chi^2(1) = 2.79; p = 0.095$ | $p = 0.076$ |
| | (Non-predictive - (NoTMS+Cz)/2) UP vs. DWN | $\chi^2(1) = 0.0011; p = 0.97$ | $p = 0.96$ |
| | (Predictive - (NoTMS+Cz)/2) UP vs. DWN | $\chi^2(1) = 0.0029; p = 0.96$ | $p = 0.96$ |
| | (Non-predictive - (NoTMS+Cz)/2) L vs. R | $\chi^2(1) = 0.79; p = 0.32$ | $p = 0.30$ |
| | (Predictive - (NoTMS+Cz)/2) L vs. R | $\chi^2(1) = 0.26; p = 0.61$ | $p = 0.61$ |
| *mIPS/IMD* | Non-predictive | $\chi^2(1) = 0.41; p = 0.52$ | $p = 0.51$ |
| | Predictive | $\chi^2(1) = 0.30; p = 0.58$ | $p = 0.57$ |
| | (Non-predictive - (NoTMS+Cz)/2) UP vs. DWN | $\chi^2(1) = 0.39; p = 0.53$ | $p = 0.52$ |
| | (Predictive - (NoTMS+Cz)/2) UP vs. DWN | $\chi^2(1) = 1.08; p = 0.30$ | $p = 0.27$ |
| | (Non-predictive - (NoTMS+Cz)/2) L vs. R | $\chi^2(1) = 0.15; p = 0.70$ | $p = 0.68$ |
| | (Predictive - (NoTMS+Cz)/2) L vs. R | $\chi^2(1) = 0.13; p = 0.72$ | $p = 0.72$ |

Note: rTMS = repetitive Transcranial Magnetic Stimulation; DV = Dependent variable; SPOC = Superior Parietal Occipital Cortex; mIPS = Medial Intraparietal Sulcus; CE<sub>x</sub> = horizontal interception error; IMD: initial movement direction; NoTMS = conditions without rTMS; Cz = conditions with rTMS applied to the control site Cz; UP = upward target motion; DWN = downward target motion; L = stimulation applied in the left hemisphere; R = stimulation applied in the right hemisphere.

**Supplementary Table 3:**
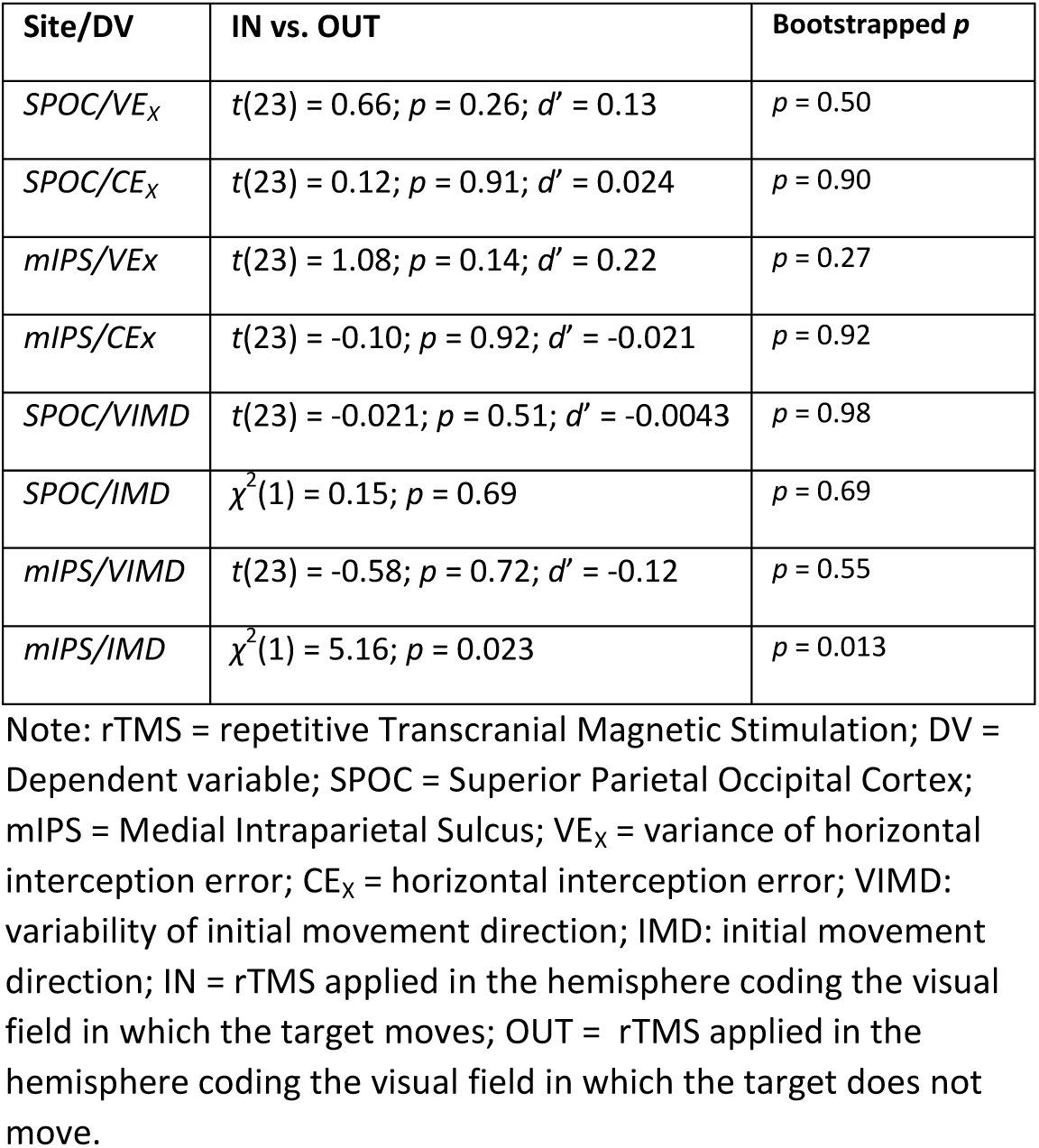
rTMS effects for non-diagonal target motion on spatial movement parameters

**Supplementary Table 4:**
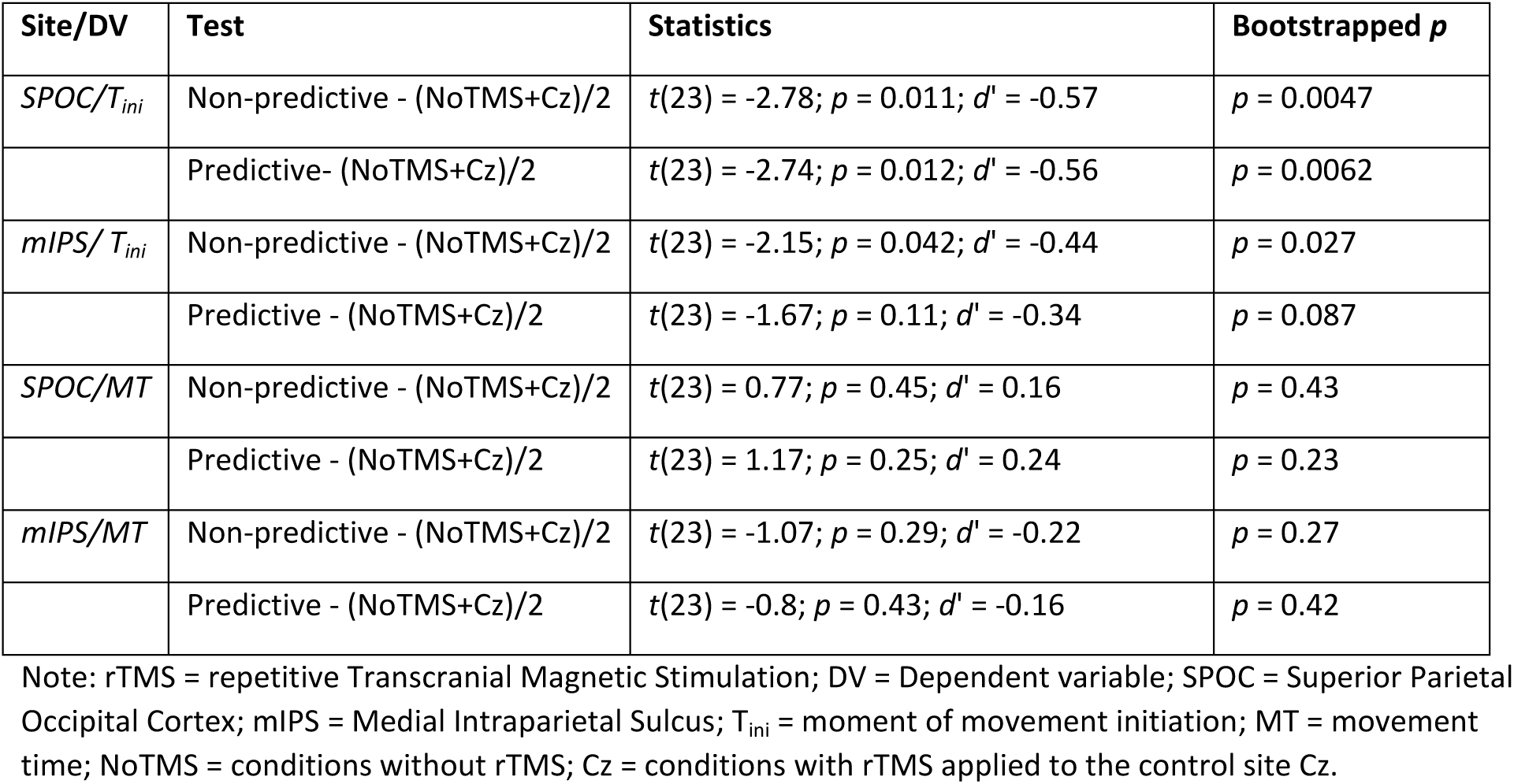
Non-predictive and Predictive rTMS effects on movement timing

| Site/DV | Test | Statistics | Bootstrapped *p* |
| --- | --- | --- | --- |
| *SPOC/T<sub>ini</sub>* | Non-predictive - (NoTMS+Cz)/2 | $t(23) = -2.78; p = 0.011; d' = -0.57$ | $p = 0.0047$ |
| | Predictive- (NoTMS+Cz)/2 | $t(23) = -2.74; p = 0.012; d' = -0.56$ | $p = 0.0062$ |
| *mIPS/T<sub>ini</sub>* | Non-predictive - (NoTMS+Cz)/2 | $t(23) = -2.15; p = 0.042; d' = -0.44$ | $p = 0.027$ |
| | Predictive - (NoTMS+Cz)/2 | $t(23) = -1.67; p = 0.11; d' = -0.34$ | $p = 0.087$ |
| *SPOC/MT* | Non-predictive - (NoTMS+Cz)/2 | $t(23) = 0.77; p = 0.45; d' = 0.16$ | $p = 0.43$ |
| | Predictive - (NoTMS+Cz)/2 | $t(23) = 1.17; p = 0.25; d' = 0.24$ | $p = 0.23$ |
| *mIPS/MT* | Non-predictive - (NoTMS+Cz)/2 | $t(23) = -1.07; p = 0.29; d' = -0.22$ | $p = 0.27$ |
| | Predictive - (NoTMS+Cz)/2 | $t(23) = -0.8; p = 0.43; d' = -0.16$ | $p = 0.42$ |

Note: rTMS = repetitive Transcranial Magnetic Stimulation; DV = Dependent variable; SPOC = Superior Parietal Occipital Cortex; mIPS = Medial Intraparietal Sulcus; $T_{ini}$ = moment of movement initiation; MT = movement time; NoTMS = conditions without rTMS; Cz = conditions with rTMS applied to the control site Cz.

**Supplementary Table 5:** rTMS effects for non-diagonal target motion on movement timing

| Site/DV | STIM<sub>IN</sub> vs. STIM<sub>OPPOSITE</sub> | Bootstrapped *p* |
| --- | --- | --- |
| *SPOC/T<sub>ini</sub>* | $t(23) = -1.91; p = 0.068; d' = -0.39$ | $p = 0.050$ |
| *SPOC/MT* | $t(23) = 1.3; p = 0.21; d' = 0.27$ | $p = 0.18$ |
| *mIPS/T<sub>ini</sub>* | $t(23) = 0.43; p = 0.67; d' = 0.088$ | $p = 0.66$ |
| *mIPS/MT* | $t(23) = -1.44; p = 0.16; d' = -0.29$ | $p = 0.14$ |

Note: rTMS = repetitive Transcranial Magnetic Stimulation; DV = Dependent variable; SPOC = Superior Parietal Occipital Cortex; mIPS = Medial Intraparietal Sulcus; T<sub>ini</sub> = moment of movement initiation; MT = movement time; STIM<sub>IN</sub> = rTMS applied in the hemisphere coding the visual field in which the target moves; STIM<sub>OPPOSITE</sub> = rTMS applied in the hemisphere coding the visual field in which the target does not move.

**Supplementary Table 6:** MNI coordinates of SPOC and mIPS in current and previous studies

|  |  | Left Hemisphere |  |  | Right Hemisphere |  |  |
| --- | --- | --- | --- | --- | --- | --- | --- |
|  | N | X | Y | Z | X | Y | Z |
| **mIPS** |  |  |  |  |  |  |  |
| This Study | 24 | -38.3 ±9 | -49.9 ±10 | 47.0 ±10 | 37.6 ±8 | -47.5 ±12 | 50.0 ±10 |
| Davare et al. (2015)<sup>15</sup> | 6 | -32 | -49 | 46 | 33 | -46 | 49 |
| Davare et al. (2012)<sup>14</sup> | 9 | -32 ±5 | -49 ±6 | 46 ±9 | 33 ±5 | -46 ±7 | 49 ±10 |
| Vesia et al. (2010)<sup>12</sup> | 6 | -22 | -68 | 46 | 25 | -61 | 46 |
| **SPOC** |  |  |  |  |  |  |  |
| This Study | 25 | -9.9 ±7 | -80.8 ±8 | 38.3 ±11 | 10.2 ±12 | -79.2 ±9 | 40.2 ±10 |
| Dessing et al. (2013)<sup>24</sup> | 7 | -6 ±1 | -81 ±2 | 35 ±6 |  |  |  |
| Vesia et al. (2010)<sup>12</sup> | 6 | -10 | -85 | 46 | 10 | -87 | 48 |

Note: mIPS = medial Intraparietal Sulcus; SPOC = Superior Parietal Occipital Cortex.

